# Genetic structure of the Coconut Rhinoceros Beetle (*Oryctes rhinoceros*) population and the incidence of its biocontrol agent (Oryctes rhinoceros nudivirus) in the South Pacific Islands

**DOI:** 10.1101/2020.07.30.229872

**Authors:** Kayvan Etebari, James Hereward, Apenisa Sailo, Emeline M. Ahoafi, Robert Tautua, Helen Tsatsia, Grahame V Jackson, Michael J. Furlong

## Abstract

Incursions of the Coconut rhinoceros beetle (CRB), *Oryctes rhinoceros*, have been detected in several countries of the south west Pacific in recent years, resulting in an expansion of the pest’s geographic range. It has been suggested that this resurgence is related to an *O. rhinoceros* mitochondrial lineage (previously referred to as the CRB-G “biotype”) that is reported to show reduced susceptibility to the well-established classical biocontrol agent, Oryctes rhinoceros nudivirus (OrNV). We investigated *O. rhinoceros* population genetics and the OrNV status of adult specimens collected in the Philippines and seven different South Pacific island countries (Fiji, New Caledonia, Papua New Guinea (PNG), Samoa, Solomon Islands, Tonga, and Vanuatu). Based on the presence of single nucleotide polymorphisms (*snp*s) in the mitochondrial Cytochrome C Oxidase subunit I (*CoxI*) gene, we found three major mitochondrial lineages (CRB-G, a PNG lineage (CRB-PNG) and the South Pacific lineage (CRB-S)) across the region. Haplotype diversity varied considerably between and within countries. The *O. rhinoceros* population in most countries was monotypic and all individuals tested belonged to a single mitochondrial lineage (Fiji, CRB-S; Tonga, CRB-S; Vanuatu, CRB-PNG; PNG (Kimbe), CRB-PNG; New Caledonia CRB-G; Philippines, CRB-G). However, in Samoa we detected CRB-S and CRB-PNG and in Solomon Islands we detected all three haplotype groups. Genotyping-by-Sequencing (GBS) methods were used to genotype 10,000 *snp*s from 230 insects across the Pacific and showed genetic differentiation in the *O. rhinoceros* nuclear genome among different geographical populations. The GBS data also provided evidence for gene flow and admixture between different haplotypes in Solomon Islands. Therefore, contrary to earlier reports, CRB-G is not solely responsible for damage to the coconut palms reported since the pest was first recorded in Solomon Islands in 2015. We also PCR-screened a fragment of OrNV from 260 insects and detected an extremely high prevalence of viral infection in all three haplotypes in the region. We conclude that the haplotype groups CRB-G, CRB-S, and PNG, do not represent biotypes, subspecies, or cryptic species, but simply represent different invasions of *O. rhinoceros* across the Pacific. This has important implications for management, especially biological control, of Coconut rhinoceros beetle in the region.

## 1. Introduction

Coconut rhinoceros beetle (CRB), *Oryctes rhinoceros* (L.) (Coleoptera: Scarabaeidae) is a serious threat to livelihoods in the Pacific islands, where coconut (*Cocos nucifera*), “the tree of life”, is an essential food, fibre and timber resource that also provides coastal protection for five million vulnerable people. *Oryctes rhinoceros* was unintentionally introduced to Samoa from its native Southeast Asia in 1909 and it has since spread to many parts of the region, causing devastating losses of coconut palms (Zelazny, 1977b). In 2015 the pest was recorded in Solomon Islands for the first time (Marshall *et al*., 2017) where it threatens the country’s SBD$140 million coconut industry and the livelihoods of 40,000 rural households (Tsatsia *et al*., 2018).

The introduction of Oryctes rhinoceros nudivirus (OrNV) from Malaysia into Samoa in the 1960s (Marschal, 1970), and then elsewhere in the region, resulted in suppression of *O. rhinoceros* populations and reduced damage to palms (Bedford 2013; Huger 2005). The biological control *O. rhinoceros* by OrNV is widely considered to be a landmark example of classical biological control and it represents one of the few examples of classical biological control involving an entomopathogen (Caltagirone *et al*., 1981). As such, the system is of considerable ecological and applied significance. New incursions of *O. rhinoceros* into countries and territories that were previously free of the pest have been reported in recent years, beginning in Guam (2007) and followed by Hawaii (2013), Solomon Islands (2015) and most recently Vanuatu and New Caledonia (2019). It has been suggested that this resurgence and spread of the pest is related to an OrNV-tolerant haplotype, “CRB-G”, which is identified by single nucleotide polymorphisms (*snp*s) in the Cytochrome C Oxidase subunit I (*CoxI*) mitochondrial gene (Marshall *et al*., 2017). This haplotype group has previously been identified throughout much of the beetle’s native and invasive ranges, including Indonesia, the Philippines, Taiwan, Palau, Guam and Solomon Islands (Marshall *et al*. 2017; Reil *et al*. 2016).

Currently, there is no genetic evidence explaining why the CRB-G haplotype might be less susceptible to OrNV than the CRB-S haplotype that first established in the region. Although, the possibility of *O. rhinoceros* tolerance/ resistance to the virus and/ or the presence of low virulence isolates of virus in different regions has been proposed (Zelazny and Alfiler 1991; Zelazny *et al*. 1992), there is no compelling evidence to support this (Huger 2005). OrNV causes chronic infection rather than rapid mortality in its adult hosts, reducing feeding, flight activity, oviposition and mating (Zelazny, 1977a, b). Such chronic effects do not necessarily cause immediate declines in adult populations and studies have shown virus introductions into new islands can decrease adult populations by as little as 10-20% of the original beetle population density (Zelazny and Alfiler 1991). The impact of OrNV can, however, be considerable, especially when it is integrated into a wider management approach.
The CRB-S and CRB-G haplotype groups only co-occur in a small number of places, and they have been referred to as biotypes. However, data to support the contention that they are biologically and/or ecologically distinct is lacking and it has not previously been established whether or not *O. rhinoceros* from the different haplotype groups interbreed freely with each other when they co-occur in sympatry. This study set out to characterize the genetic variability of *O. rhinoceros* populations sampled across the Pacific Islands and to estimate the incidence of OrNV infection in the sampled insects. By combining a genotyping-by-sequencing approach with *CoxI* sequencing we were able to investigate gene flow across the different haplotype groups using multiple nuclear *snp* markers.

## 2. Materials and Methods

### 2.1 Insect Collection

Adult *O. rhinoceros* were collected in traps baited with aggregation pheromone (Oryctalure, P046-Lure, ChemTica Internacional, S. A., Heredia, Costa Rica) lures. Collections were made in Fiji (Viti Levu and Vanua Levu), New Caledonia (Nouméa), Papua New Guinea (Kimbe, New Britain), Samoa (Upolu and Savai’i), Solomon Islands (Guadalcanal, Russel Islands, Gizo, Kolombangara and Santa Cruz Islands), Tonga (Tongatapu), and Vanuatu (Efate) in the south west Pacific and in the Philippines (Los Banos) between January and October, 2019. Upon removal from traps, individual beetles were preserved in 95% ethanol in separate containers and shipped to the University of Queensland, Brisbane, Australia for further analysis.

### 2.2 DNA extraction and mitochondrial *CoxI* gene analysis

To characterise the different mitochondrial DNA haplotypes in the *O. rhinoceros* samples from across the region, we sequenced the *CoxI* gene in 260 individuals. DNA was extracted from a leg of adult *O. rhinoceros* using a DIY spin column protocol using 96-well silica filter plates from Epoch life sciences (Missouri City, TX, USA) (Ridley *et al*. 2016). A small fragment of the *CoxI* gene was amplified in these individuals with the primers LCO1490 and HCO2198 (Folmer *et al*. 1994). PCRs were carried out as 25 µl reactions containing 0.03 units of MyTaq™ (Bioline), 1 × buffer and 0.2 μM of forward and reverse primers and 2 μl of template DNA. The PCR conditions were as follows: 95°C for 3 min, 40 cycles consisting of denaturation at 95°C for 15s, annealing at 55°C for 30s and 72°C for 15s. PCR products were cleaned using one unit each of Exonuclease I and Antarctic Phosphatase (New England Biolabs, Massachusetts, USA), with sequencing reactions performed using an ABI3730 Genetic Analyzer (Applied Biosystems) at Macrogen Inc. Seoul, South Korea. PCR products were sequenced bi-directionally after confirmation by 1% agarose gel electrophoresis. After primer trimming and alignment, these two *CoxI* regions overlapped by 621 bp and this segment was used for further phylogenetic analysis using CLC genomic workbench version 12. We used the Jukes-Cantor algorithm to measure nucleotide distance under the UPGMA model and ran multiple gene alignment for overlapped regions among the samples from different countries. The haplotype network was generated based on 621 bp of the *CoxI* gene region from a total of 260 individuals by PopART 1.7.2 (Leigh and Bryant 2015).

#### Genotyping-by-Sequencing analysis

To assess gene flow across the nuclear genome of these populations we generated genome-wide *snp* data using a genotyping-by-sequencing method. We followed the protocol and adaptor regime described by (Hereward *et al*. 2020), which is based on the methods of (Elshire *et al*. 2011), (Poland *et al*. 2012) (Peterson *et al*. 2012), with barcodes based on (Caporaso *et al*. 2012). Briefly, samples were normalised to 200ng and double-digested with MspI and PstI, we ligated 96 unique forward barcodes, and three different plates of reverse barcodes so that every sample had a unique forward and reverse barcode combination. Each plate of 96 samples was then pooled and size selected on a Blue Pippin (Sage Science) to 300-400bp. Each of these pools was PCR amplified to complete the adaptors and add indices. We then normalised and pooled these three PCR products for a total of 288 individuals per sequencing lane, and sequenced the libraries with PE150 Illumina sequencing at Novogene (Beijing, China).

The sequence data were demultiplexed, assembled, and *snps* called using STACKS (Catchen *et al* 2013). We then filtered the vcf using vcftools, with the data first being filtered to a minor allele count of three (one heterozygote and one homozygote), which allows the conservative removal of singleton *snps* that are likely to be errors, without discarding rare alleles (Linck & Battey 2019). We set a minimum depth of 5, and kept only biallelic *snps*. We filtered the data for missing data in three steps. First, any marker missing more than 50% data was discarded (i.e. 50% of individuals were not genotyped at that marker), to remove the markers most affected by missing data. Second, any individual that had missing data at more than 50% of the markers was discarded, to remove the individuals that had poor quality genotyping. Finally, any marker missing more than 5% data was discarded to produce a filtered dataset with relatively little missing data (∼3%). We plotted a PCA (Principal Component Analysis) graph using adegenet package in R (Jombart 2008), and then ran *structure* (Pritchard *et al*. 2000). *Structure* is an individual-based clustering algorithm that assigns individuals to each of *K* population clusters using Hardy-Weinberg equilibrium and linkage information in genetic markers. We kept only one *snp* per locus, so the total number of *snps* was reduced to 9,609.

#### Incidence of OrNV infection in adult *O. rhinoceros*

To determine the incidence of OrNV in *O. rhinoceros* specimens collected in the field from across the Pacific, the gut tissue was carefully dissected from adult beetles and total DNA from individual specimens was extracted as described previously. The presence of OrNV in gut tissue was confirmed by PCR amplification of a 945 bp product using the OrV15 primers that target the OrNV-gp083 gene (Richards *et al*. 1999). The PCR product was visualized by gel electrophoresis and validated by Sanger sequencing.

## Results

### Mitochondrial *CoxI* analysis

Twelve different haplotypes, each belonging to one of three major mitochondrial lineages (CRB-S, CRB-G and CRB-PNG) were identified in field-collected *O. rhinoceros* (Fig. 1). These sequences have been submitted to NCBI under GenBank accession numbers MN809502-MN809525.

Haplotype diversity varied considerably between regions, and between and within countries. In Tonga, the population was completely monotypic and all individuals tested belonged to a single mitochondrial lineage (CRB-S; Fig. 1). The population in Fiji contained four different haplotypes, all of which also belonged to the CRB-S lineage (Fig. 1). In samples from Samoa, three distinct haplotypes were identified; two of these were more frequent in the population and both belonged to the CRB-S lineage. A third haplotype, which differed from the other haplotypes in Samoa by 4 to 6 base pairs (Figs.1 and 2; Table 1), belonged to the CRB-PNG lineage. The haplotype network generated for the *CoxI* gene suggests that *O. rhinoceros* from Tonga and Fiji, and most of those collected from Samoa, are almost identical with low mitochondrial genetic variation between these individuals (Fig. 2 and Table 1).

**Table 1.** Pairwise comparison of *CoxI* gene sequence for 12 identified CRB haplotypes in the Pacific. Upper comparisons represent the number of single nucleotide differences between haplotypes and lower comparisons are the percentage of similarity between haplotypes.

| Mitochondrial Lineages |  | Haplotype #1 | Haplotype #2 | Haplotype #3 | Haplotype #4 | Haplotype #5 | Haplotype #6 | Haplotype #7 | Haplotype #8 | Haplotype #9 | Haplotype #10 | Haplotype #11 | Haplotype #12 |
| --- | --- | --- | --- | --- | --- | --- | --- | --- | --- | --- | --- | --- | --- |
| CRB-PNG | Haplotype #1 |  | 1 | 1 | 7 | 5 | 6 | 6 | 6 | 7 | 8 | 6 | 8 |
| CRB-PNG | Haplotype #2 | 99.84 |  | 2 | 6 | 4 | 5 | 5 | 5 | 6 | 7 | 5 | 7 |
| CRB-PNG | Haplotype #3 | 99.84 | 99.68 |  | 6 | 4 | 5 | 5 | 5 | 6 | 7 | 5 | 7 |
| CRB-S | Haplotype #4 | 98.87 | 99.03 | 99.03 |  | 2 | 3 | 3 | 3 | 4 | 5 | 5 | 3 |
| CRB-S | Haplotype #5 | 99.19 | 99.36 | 99.36 | 99.68 |  | 1 | 1 | 1 | 2 | 3 | 3 | 3 |
| CRB-S | Haplotype #6 | 99.03 | 99.19 | 99.19 | 99.52 | 99.84 |  | 2 | 2 | 3 | 4 | 4 | 4 |
| CRB-S | Haplotype #7 | 99.03 | 99.19 | 99.19 | 99.52 | 99.84 | 99.68 |  | 2 | 3 | 4 | 4 | 4 |
| CRB-S | Haplotype #8 | 99.03 | 99.19 | 99.19 | 99.52 | 99.84 | 99.68 | 99.68 |  | 3 | 4 | 4 | 4 |
| CRB-G | Haplotype #9 | 98.87 | 99.03 | 99.03 | 99.36 | 99.68 | 99.52 | 99.52 | 99.52 |  | 1 | 1 | 1 |
| CRB-G | Haplotype #10 | 98.71 | 98.87 | 98.87 | 99.19 | 99.52 | 99.36 | 99.36 | 99.36 | 99.84 |  | 2 | 2 |
| CRB-G | Haplotype #11 | 99.03 | 99.19 | 99.19 | 99.19 | 99.52 | 99.36 | 99.36 | 99.36 | 99.84 | 99.68 |  | 2 |
| CRB-G | Haplotype #12 | 98.71 | 98.87 | 98.87 | 99.52 | 99.52 | 99.36 | 99.36 | 99.36 | 99.84 | 99.68 | 99.68 |  |

Haplotype #1: Solomon Islands (Malo; Santa Cruz, Gizo, Guadalcanal and Russell Island), PNG (New Britain) and Vanuatu
Haplotype #2: Samoa
Haplotype #3: Solomon Islands (Guadalcanal)
Haplotype #4: Samoa
Haplotype #5: Fiji, Tonga, Samoa and Solomon Islands (Guadalcanal)
Haplotype #6: Fiji
Haplotype #7: Fiji
Haplotype #8: Fiji
Haplotype #9: Solomon Islands (Guadalcanal, Kolombangara; Russell Island), Philippines and New Caledonia

All *O. rhinoceros* individuals collected from PNG (Kimbe, New Britain) belonged to the CRB-PNG haplotype lineage and insects conforming to this haplotype lineage were also identified in samples from Vanuatu, and Solomon Islands (Malo, Santa Cruz Islands; Gizo; Guadalcanal and Russell Islands) (Figs. 1 and 2; Table 1). Insects with a high degree of similarity to this haplotype group were identified from Solomon Islands (Guadalcanal) and Samoa (Figs. 1 and 2). This haplotype group showed maximum polymorphism (5-8 nucleotide differences) compared to other haplotypes collected from the south Pacific islands and the Philippines (Table 1).

Four haplotypes conforming to the haplotype group previously described as CRB-G were detected in insects collected in Solomon Islands (Guadalcanal, Russel Islands and Kolombangara), New Caledonia, and in the Philippines (Figs. 1 and 2; Table 1). This group of insects showed more similarity to the CRB-S lineage than to the CRB-PNG lineage based on the number of SNPs in their *CoxI* gene (Table 1).

**Figure 1.**
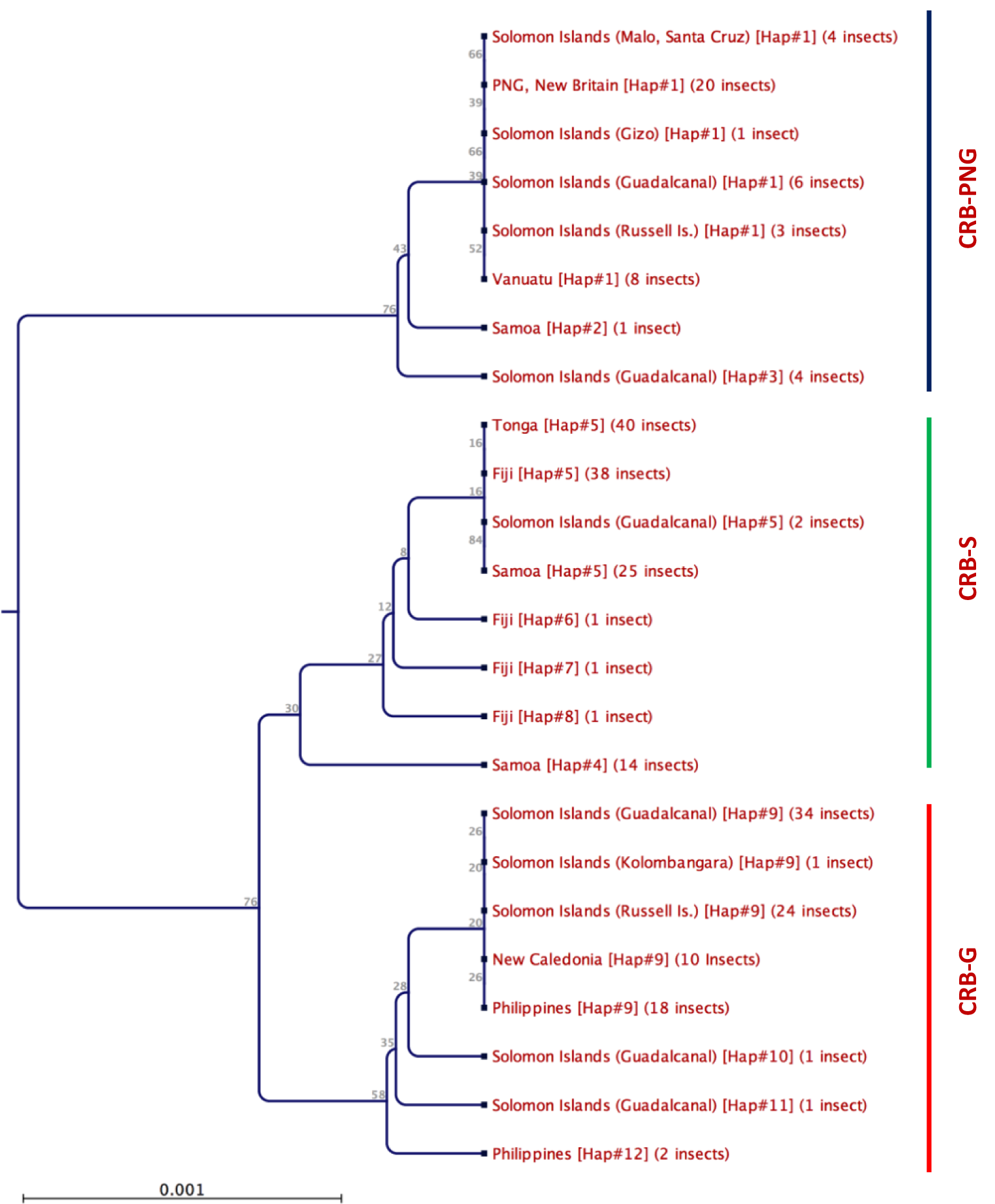
Phylogeny of *O. rhinoceros* based on partial *CoxI* gene sequence (621bp). 12 different haplotypes of *O. rhinoceros*, in three major haplotype clades (CRB-PNG, CRB-S and CRB-G) were identified.

**Figure 2.**
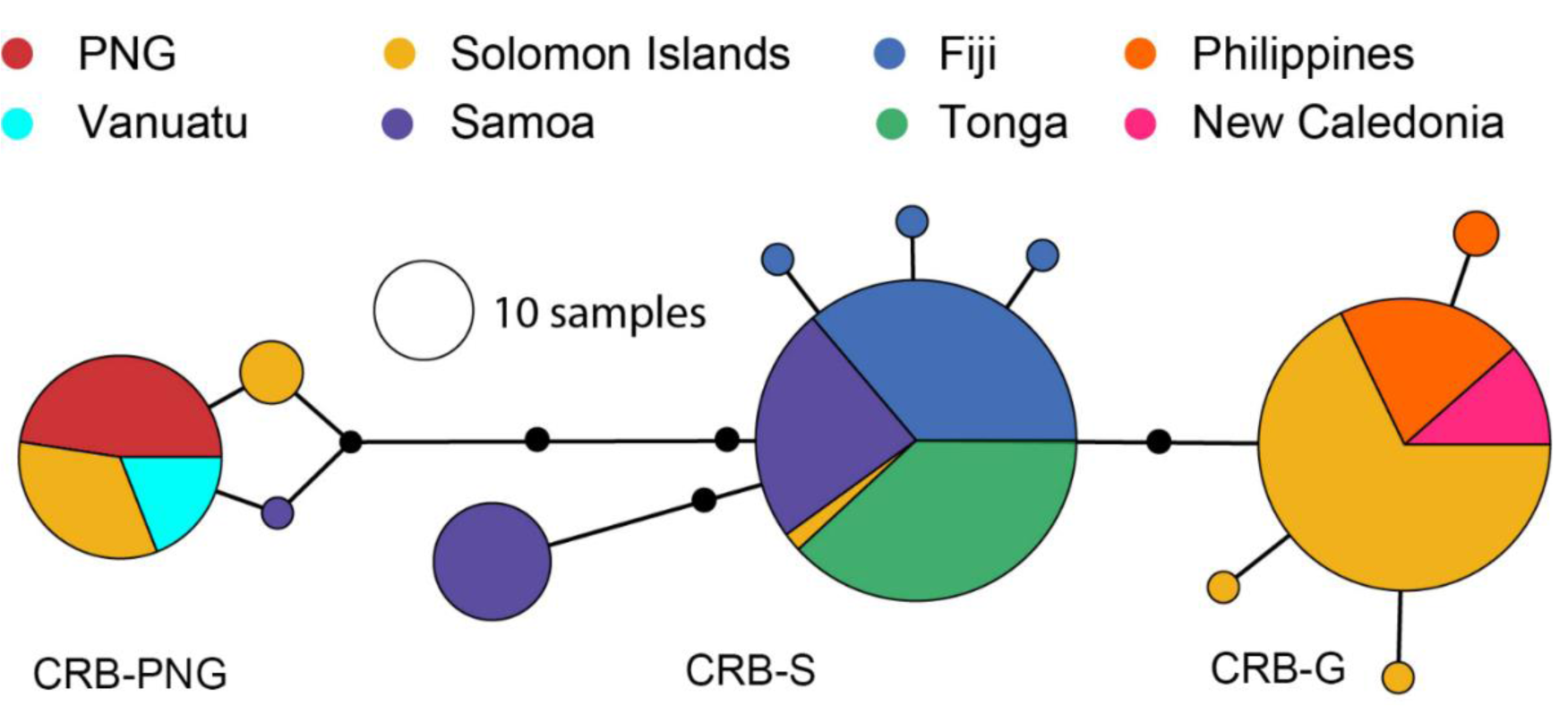
*O. rhinoceros CoxI* haplotype network. The network is based on 621 bp of the *CoxI* gene region from a total of 260 individuals collected from Pacific Islands (created in PopART (Leigh and Bryant 2015)). The circles represent haplotypes and their diameters are proportional to relative abundance across all samples in the study (black circles along branches represent single nucleotide differences). The different colours show the distribution of the different haplotypes between the different countries.

### Genotyping-by-Sequencing analysis

Data based on nuclear DNA analysis shows the *O. rhinoceros* population in Solomon Islands is admixed, with both the CRB-G and CRB-PNG haplotypes showing clear evidence of gene flow in the PCA plot and in the structure analysis (Fig. 3). Specimens from Tonga, Fiji and Samoa clustered very close to each other in the PCA, with samples from Tonga and Fiji clustering particularly close together (Fig. 3A). The beetles from all three of these countries belong to the CRB-S mitochondrial haplotype group (Fig 3B), and the genetic differentiation detected between them has likely occurred since they initially invaded the region. The beetles collected from PNG clustered close together in the PCA (Fig. 3A) and are distinct from those collected in Solomon Islands, despite the admixture between the CRB-PNG and CRB-G mitochondrial lineages there (Fig. 3B). The STRUCTURE analysis with K=4 had the best support based on the delta K method, and this placed PNG and Solomon Islands together (likely due to the recent admixture), but separated Fiji, Tonga and Samoa into different clusters (Fig 3C). We only collected two individuals from the CRB-S mitochondrial lineage from Guadalcanal in this study and we were unable to include them in our GBS analysis at the time of investigation.

**Figure 3.**
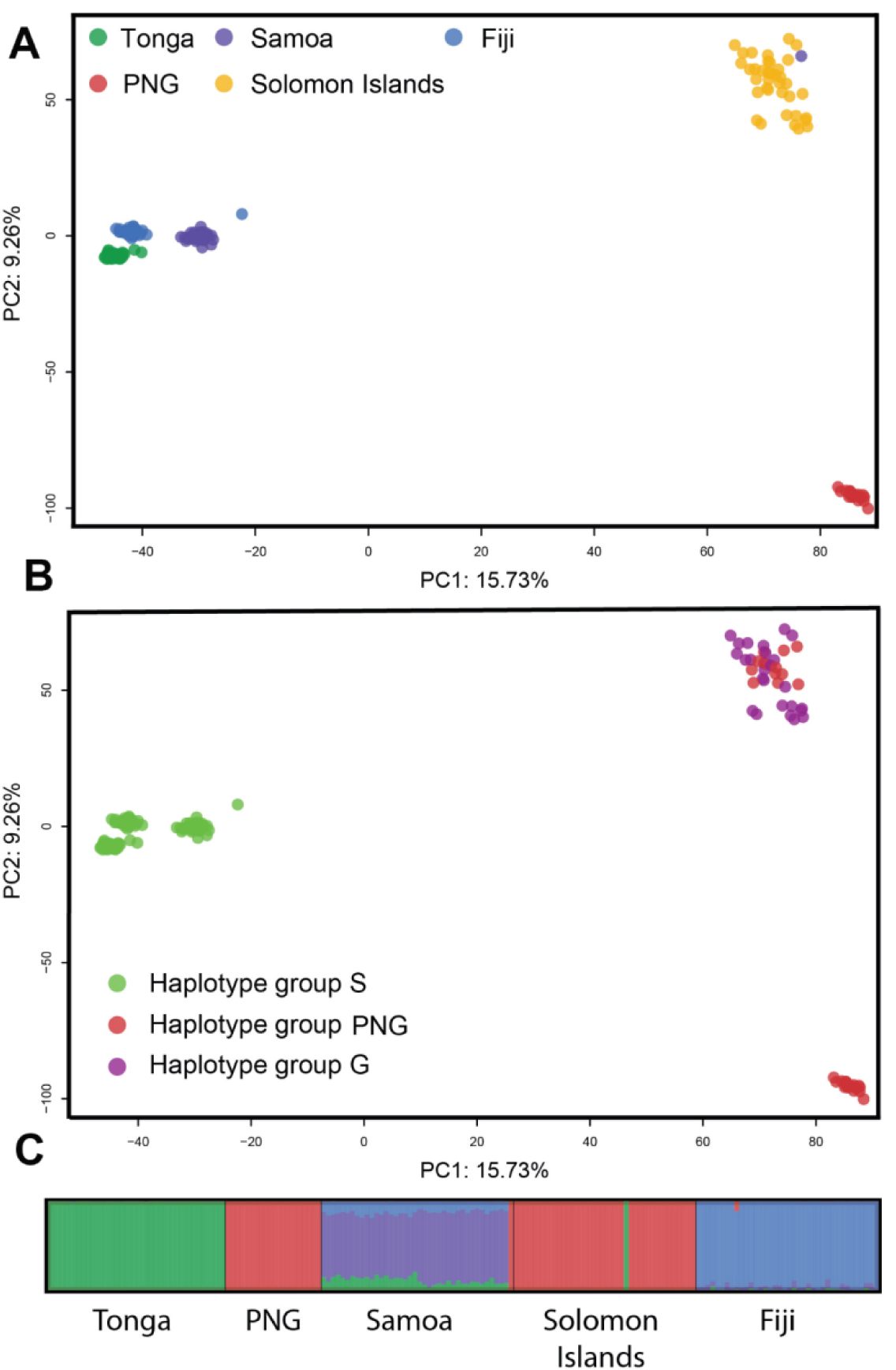
PCA plots of the GBS data coloured by country of origin (A), and by *CoxI* haplotype lineage (B). A STRUCTURE plot (C) shows the assignment of individuals to each of four genetic clusters, each bar represents one individual and the colour indicates the posterior probability of assignment to each cluster

### Incidence of OrNV infection in adult *O. rhinoceros*

OrNV infection was detected in adult *O. rhinoceros* in all countries and infection rates were extremely high (63-95% where ≥ 10 individual insects were tested) in all mitochondrial lineages (Table 2). We detected several OrNV infected beetles from each of the different haplotypes collected in Solomon Islands. Detection of OrNV infection in the small number of specimens collected from more remote area like the Santa Cruz Islands demonstrates that OrNV is widely distributed in *O. rhinoceros* populations in Solomon Islands, but it does not provide reliable information on the prevalence of the pathogen in those outlaying region. Nonetheless, testing 96 wild caught *O. rhinoceros* adults collected in Guadalcanal and Russel Islands indicated that >80% of beetles were infected with OrNV and similarly high infection rates (>90%) were recorded in *O. rhinoceros* collected from New Caledonia and Vanuatu (Table 2).

**Table 2.** Incidence of OrNV infection in Adult *O. rhinoceros*.

| *O. rhinoceros* origin |  | *O. rhinoceros* mitochondrial lineage |  |  |  |  |  | Overall infection rate (%) |
| --- | --- | --- | --- | --- | --- | --- | --- | --- |
| Country | Location | CRB-S |  | CRB-PNG |  | CRB-G |  |  |
|  |  | No. tested | No. OrNV +ve (%) | No. tested | No. OrNV +ve (%) | No. tested | No. OrNV +ve (%) |  |
| Fiji | Viti Levu | 21 | 21 (100) | - | - | - | - | 100 |
|  | Vanua Levu | 17 | 15 (88) | - | - | - | - | 88 |
| Solomon Islands | Guadalcanal, Dorma | 2 | 0 (0) | 12 | 9 (75) | 26 | 16 (64) | 63 |
|  | Guadalcanal, Mbalasuna | - | - | 17 | 17 (100) | 21 | 21 (100) | 100 |
|  | Russel Islands | - | - | - | - | 18 | 16 (89) | 89 |
|  | Gizo | - | - | 1 | 0 (0) | - | - | 0 |
|  | Kolombangara | - | - | - | - | 1 | 0 (0) | 0 |
|  | Santa Cruz Islands | - | - | 4 | 2 (50) | - | - | 50 |
| Tonga | Nuku'alofa | 20 | 19 (95) | - | - | - | - | 95 |
| Samoa | Upolu | 10 | 7 (70) | 1 | 1 (100) | - | - | 73 |
|  | Savai'i | 18 | 18 (100) | - | - | - | - | 100 |
| New Caledonia | Noumea | - | - | - | - | 10 | 10 (100) | 100 |
| Vanuatu | Efate | - | - | 5 | 4 (80) | - | - | 80 |
| Papua New Guinea | New Britain, Kimbe | - | - | 15 | 11 (73) | - | - | 73 |
| Philippines | Los Baños | - | - | - | - | 9 | 7 (78) | 78 |

## Discussion

Understanding gene flow and genetic variation in *O. rhinoceros* populations can help explain active evolutionary processes occurring within the species and contribute to improved management strategies. Previously, Reil *et al*. (2016) examined the *CoxI* and carbamoyl-phosphate synthetase, aspartate transcarbamylase, dihydroorotase (*Cad* gene) nuclear genes in *O. rhinoceros* collected from their native range in Asia and the invaded islands of Palau, Oahu, Guam and American Samoa in the Pacific, but lack of meaningful variation in the *CoxI* and *Cad* genes prevented them from generating clear evidence for *O. rhinoceros* movement. More recently, Marshall *et al*. (2017) suggested that two major mitochondrial *CoxI* haplotype groups correspond to two invasion fronts of *O. rhinoceros* across the Pacific and that one of them, the CRB-G haplotype group, was associated with increased tolerance to resistance to OrNV. These haplotype groups have been referred to as “biotypes”, but it is not clear if any biological features of the insect are tied to these two mitochondrial lineages.

In this study we combined multiple nuclear markers with the mitochondrial *CoxI* data and OrNV screening to provide clarity on the genetic structure of the *O. rhinoceros* population across the south Pacific and on the relationships between the different haplotype groups and OrNV infection. The *O. rhinoceros* population in most countries was monotypic, with all individuals tested belonging to a single mitochondrial lineage; in Fiji and Tonga insects were from the CRB-S haplotype group, in PNG and Vanuatu they were from the CRB-PNG group and in New Caledonia they were from the CRB-G group (Fig. 1). However, in Samoa insects from both the CRB-S and CRB-PNG haplotype groups were detected and in Solomon Islands all three haplotype groups were represented in samples (Fig. 1). While the incidence of OrNV infection in samples varied, high levels of infection were recorded in all *O. rhinoceros* haplotypes wherever they were found (Table 2).

In Solomon Islands, admixture and gene flow between CRB-PNG and CRB-G was detected when both mitochondrial lineages were present in the same geographical location. Our also data shows that the CRB-PNG haplotype is more distantly related to the CRB-G haplotype than is the CRB-S haplotype, suggesting that CRB-S and CRB-G haplotypes are also likely to become admixed when they are brought into sympatry. Such a pattern has been observed in Palau where both the CRB-S and the CRB-G haplotypes are established (Kitalong *et al*., 2018; Reil *et al*., 2018). These different haplotype groups therefore do not represent cryptic species, subspecies, or biotypes; more simply, they represent separate invasion fronts of *O. rhinoceros* across the Pacific.

The current *CoxI* based haplotype diagnostics likely do not reflect any significant biological differences between insects and they are only diagnostic for movement of beetles of a given haplotype lineage into a region. The lack of meaningful mitochondrial diversity in *O. rhinoceros* populations from Tonga, Samoa and Fiji probably reflects a shared invasion history in these countries. In places such as Solomon Islands, where multiple haplotype groups are present and there is obvious introgression between them (Fig. 3), the utility of *CoxI* based haplotype diagnostics in even this respect is limited as insects can only be traced back through the maternal line. In areas of introgression, they cannot be used to draw meaningful conclusions about the movement of insects or their likely susceptibility to OrNV. For example, male insects from the CRB-G mitochondrial lineage that mate with female insects from a different haplotype would produce offspring undetectable as CRB-G using *CoxI* haplotype diagnostics, but they would carry paternal nuclear genes, including those mediating interations with viruses and other pathogens. Similarly, females from the CRB-G mitochondrial lineage that mate with males from another haplotype would produce offspring detectable as CRB-G but the genetic material in their nuclear genome would be from both parents.

Misinterpretation of the current pest situation and the critical genetic relationships between populations can mislead our primary understanding of *O. rhinoceros* biology, waste diagnostic resources, and prevent the development of appropriate control strategies. It has been suggested that *O. rhinoceros* belonging to the CRB-G lineage cause more damage to host palms than insects from the other lineages (Marshall *et al*., 2018), but definitive evidence is lacking. Furthermore, the *O. rhinoceros* that have recently invaded Vanuatu belong to the CRB-PNG mitochondrial lineage and they are positive for OrNV, while those that have invaded New Caledonia belong to the CRB-G mitochondrial lineage and they are also positive for OrNV. This indicates that genetic and phenotypic characteristics that have been associated with CRB-G are not unique to this haplotype. Invasions of islands by *O. rhinoceros* are likely to be human mediated (Reil *et al*., 2016) and the CRB-PNG group is as able as the CRB-G group take advantage of this to expand its range, and both lineages are susceptible to OrNV. As in the Solomon Islands, individuals responsible for these new invasions could be the progeny of parents from different mitochondrial lineages (CRB-PNG and CRB-G) but they are only identifiable based on their maternal mitochondrial lineage.

Resistance to OrNV in the CRB-G mitochondrial lineage has been suggested because of an absence of OrNV infection in wild caught insects from this haplotype group and the results from laboratory bioassays investigating the susceptibility of adult *O. rhinoceros* from the Guam population of this lineage to OrNV (Marshall *et. al*., 2017). Contrary to our findings, OrNV infection was not detected in *O. rhinoceros* from the CRB-G mitochondrial lineage collected in Solomon Islands or the Philippines, but it was detected in insects from this lineage collected from Palau (Marshall *et al*. 2017), where the CRB-G and CRB-S lineages occur in sympatry. Interestingly, Kitalong *et al*., (2018) found very high rates of OrNV infection both haplotypes in Palau (83% in CRB-G and 92% in CRB-S), supporting our findings of a high incidence of OrNV infection in all haplotypes in the Pacific Islands and the Philippines (Table 2). In laboratory bioassays, oral treatment of CRB-G adults with a range of OrNV isolates failed to induce significantly greater mortality than in untreated control insects but injection of these isolates into the adult haemocoel did cause increased mortality (Marshall *et al*., 2017). Although OrNV infection can be fatal to all developmental stages *O. rhinceros* except eggs, in adults the disease is chronic, reducing feeding, flight, oviposition and male mating activity (Zelazny 1977ab) and it does not typically present with any obvious external symptoms (Burand, 1998). Such effects are typical of nudivirus infections as the viruses usually replicate in reproductive tissues and cause sterility in the host (e.g. Unckless 2011). Thus, measuring adult mortality alone is not an appropriate method to assess the pathogenicity or virulence of OrNV in its host and results of such studies are only meaningful when control insects known to be susceptible to the pathogen are included in simultaneous assays.

Low nudivirus mortality rates in infected insects may not be related to host resistance to the pathogen but rather due to changes in the virus (Hill and Unckless, 2011). We recently determined the full length of the OrNV genomic sequence from an infected *O. rhinoceros* adult (CRB-G mitochondrial lineage) collected in the Solomon Islands (Etebari *et al*. 2020a). The complete circular genome of the virus has 138 amino acid modifications in 53 coding regions when compared with the originally described full genome sequence of the Ma07 strain from Malaysia (NC_011588). Although this demonstrates considerable change in the pathogen, further sequence analyses of more OrNV isolates and investigation of their effects on host insects from multiple populations is needed to determine if such mutations compromise the efficacy of OrNV as a biocontrol agent.

The origin of the OrNV that we found infecting *O. rhinoceros* in Solomon Islands is not known. Analysis of OrNV genomic structural and transcriptional variation in samples collected across the Pacific and the Philippines showed that OrNV strains from Solomon Islands and the Philippines are closely related, while those from PNG and Fiji formed a distinct adjacent clade (Etebari *et al*., 2020b). The high level of similarity between the OrNV in Solomon Islands and the Philippines and lack of evidence that OrNV from the Philippines has been introduced, suggests that the OrNV now infecting *O. rhinoceros* in Solomon Islands arrived with an incursion of infected *O. rhinoceros* that originated in Southeast Asia.

In conclusion, the different haplotypes of coconut rhinoceros beetle reported in the south west Pacific do not represent biotypes, subspecies, or cryptic species. Rather, they represent the different invasion histories of the pest. This renders *CoxI* inappropriate as a diagnostic marker for other traits, especially in countries, such as Solomon Islands, where the lineages associated with different invasion fronts have come back into sympatry and are admixed. Although we found OrNV infecting all mitochondrial lineages of *O. rhinoceros* across the Pacific, our knowledge of the host-pathogen interaction remains limited. Molecular analysis of different haplotypes of *O. rhinoceros* in response to OrNV is necessary to identify the mechanism of potential resistance to OrNV. Further work is also required to establish the evolutionary history of OrNV in these regions in combination with a better understanding of the population genetics of *O. rhinoceros* across the Pacific.

## Acknowledgements

This project was supported by the Australian Centre for International Agricultural Research funding (HORT/2016/185) and the University of Queensland (UQECR2057321). We thank Nitya N. Singh (Government of Fiji), Marie Joy Beltran (University of the Philippines Los Baños), Aurélie Chan (Department of Veterinary, Food and Rural Affairs, Government of New Caledonia) and colleagues from Papua New Guinea Oil Palm Research Association, Dami research station for assistance in collecting and providing the insect specimens for genetic analysis.

## Data Accessibility

Raw genotyping by sequencing data is accessible by NCBI accession PRJNA648153

## Authors’ contributions

Kayvan Etebari: Conceptualization, Methodology, Investigation, Data curation, Formal analysis, Visualization, Writing-Original draft preparation, Funding acquisition

James Hereward: Methodology, Investigation, Data curation, Visualization, Writing-Reviewing and Editing.

Apensia Sailo: Resources

Robert Tautua: Resources

Emeline M. Ahoafi: Resources

Helen Tsatsia: Resources

Grahame Jackson: Resources

Michael J. Furlong: Conceptualization, Investigation, Writing - Reviewing and Editing, Funding acquisition, Project administration, Supervision.

## Reference

Bedford GO (2013) Biology and Management of Palm Dynastid Beetles: Recent Advances. In: Berenbaum MR (ed) Annual Review of Entomology, Vol 58, vol 58. Annual Review of Entomology. Annual Reviews, Palo Alto, pp 353–372.

Bedford GO (2018) Possibility of evolution in culture of the Oryctes Nudivirus of the Coconut Rhinoceros Beetle *Oryctes rhinoceros* (Coleoptera: Scarabaeidae: Dynastinae). Advances in Entomology 6:27–33.

Burand, J.P., 1998. Nudiviruses. In: Miller, L.K., Ball, L.A. (Eds.), The Insect Viruses. Springer US, Boston, MA, pp. 69–90.

Caltagirone, L.E., 1981. Landmark Examples in Classical Biological Control. Annual Review of Entomology 26, 213–232.

Caporaso JG et al. (2012) Ultra-high-throughput microbial community analysis on the Illumina HiSeq and MiSeq platforms. ISME Journal 6:1621–1624.

Catchen J, Hohenlohe PA, Bassham S, Amores A, Cresko WA (2013) Stacks: an analysis tool set for population genomics. Molecular Ecology 22:3124–3140.

Crawford AM (1981) Attempts to obtain Oryctes baculovirus replication in three insect cell cultures Virology 112:625–633

Demirbas-Uzel G, Parker AG, Vreysen MJB, Mach RL, Bouyer J, Takac P, Abd-Alla AMM (2018) Impact of *Glossina pallidipes* salivary gland hypertrophy virus (GpSGHV) on a heterologous tsetse fly host, *Glossina fuscipes fuscipes* BMC Microbiol 18:12 e161

Elshire RJ, Glaubitz JC, Sun Q, Poland JA, Kawamoto K, Buckler ES, Mitchell SE (2011) A robust, simple Genotyping-by-Sequencing (GBS) approach for high diversity species. PLoS One 6 e19379

Etebari K, Filipovic I, Rasic G, Devine GJ, Tsatsia H, Furlong MJ (2020a) Complete genome sequence of Oryctes rhinoceros nudivirus isolated from the coconut rhinoceros beetle in Solomon Islands. Virus research 278:197864–197864.

Etebari, K, Parry, R, Beltran, MJB, and MJ, Furlong (2020b) Genomic structural and transcriptional variation of Oryctes rhinoceros nudivirus (OrNV) in Coconut Rhinoceros Beetle. bioRxiv 2020.05.27.119867; doi: https://doi.org/10.1101/2020.05.27.119867

Folmer O, Black M, Hoeh W, Lutz R, Vrijenhoek R (1994) DNA primers for amplification of mitochondrial cytochrome c oxidase subunit I from diverse metazoan invertebrates. Molecular Marine Biology and Biotechnology 3:294–299

Gopal M, Gupta A, Sathiamma B, Nair CPR (2002) Microbial pathogens of the coconut pest Oryctes rhinoceros: influence of weather factors on their infectivity and study of their coincidental ecology in Kerala, India. World Journal of Microbiology & Biotechnology 18:417–421.

Hereward JP, Smith TJ, Brookes DR, Gloag R, Walter GH (2020) Tests of hybridisation in *Tetragonula* stingless bees using multiple genetic markers bioRxiv:2020.2003.2008.982546 doi: 10.1101/2020.03.08.982546

Huger AM (2005) The *Oryctes* virus: Its detection, identification, and implementation in biological control of the coconut palm rhinoceros beetle, *Oryctes rhinoceros* (Coleoptera : Scarabaeidae). Journal of Invertebrate Pathology 89:78–84.

Jombart T (2008) Adegenet: a R package for the multivariate analysis of genetic markers Bioinformatics 24:1403–1405.

Kitalong, C., Ramarui, J.O., Ngiramengior, J. Skey, B., Masang, N., Watanabe, S., Melzer, M., Nakai, M. amd J Miles (2018) CRB damage and resistance assessment in the Palau Archipelago. The 2018 International Congress of Invertebrate Pathology and Microbial Control and the 51st Annual Meeting of the Society for Invertebrate Pathology, Gold Coast, Australia 12-6 August 2018. Page 48.

Leigh JW, Bryant D (2015) POPART: full-feature software for haplotype network construction Methods in Ecology and Evolution 6:1110–1116.

Linck, E, and Battey, CJ (2019) Minor allele frequency thresholds strongly affect population structure inference with genomic data sets. Molecular Ecology Resources 19 (3): 639–47

Marschal K (1970) Introduction of a new virus disease of the Coconut Rhinoceros Beetle in Western Samoa. Nature 225: 288–289.

Marshall SDG, Moore A, Vaqalo M, Noble A, Jackson TA (2017) A new haplotype of the coconut rhinoceros beetle, *Oryctes rhinoceros*, has escaped biological control by *Oryctes rhinoceros* nudivirus and is invading Pacific Islands. Journal of Invertebrate Pathology 149: 127–134.

Marshall SDG, Moore, A, Ero, M, Fanai, C, Vaqalo, M, and TA Jackson (2018) Progress with control of a virus resistant coconut rhinoceros beetle. The 2018 International Congress of Invertebrate Pathology and Microbial Control and the 51st Annual Meeting of the Society for Invertebrate Pathology, Gold Coast, Australia 12-6 August 2018. Page 47.

Moslim R, Kamarudin N, Ghani IA, Wahid MB, Jackson TA, Tey CC, Ahdly M (2011) Molecular approaches in the assessment of *Oryctes rhinoceros* virus for the control of rhinoceros beetle in oil palm plantations. Journal of Oil Palm Research 23: 1096–1109

Palmer WH, Medd NC, Beard PM, Obbard DJ (2018) Isolation of a natural DNA virus of *Drosophila melanogaster*, and characterisation of host resistance and immune responses PLoS Pathog 14:26 e1007050

Peterson BK, Weber JN, Kay EH, Fisher HS, Hoekstra HE (2012) Double digest RADseq: An inexpensive method for *de novo snp* discovery and genotyping in model and non-model Species. PLoS One 7: e37135

Poland JA, Brown PJ, Sorrells ME, Jannink JL (2012) Development of high-density genetic maps for barley and wheat using a novel two-enzyme Genotyping-by-Sequencing approach. PLoS One 7 e32253

Pritchard JK, Stephens M, Donnelly P (2000) Inference of population structure using multilocus genotype data. Genetics 155: 945–959

Pushparajan C, Claus JD, Marshall SDG, Visnovsky G (2013) Characterization of growth and Oryctes rhinoceros nudivirus production in attached cultures of the DSIR-HA-1179 coleopteran insect cell line. Cytotechnology 65:1003–1016.

Reid S, Chan LCL, Matindoost L, Pushparajan C, Visnovsky G (2016) Cell Culture for Production of Insecticidal Viruses. In: Glare TR, MoranDiez ME (eds) Microbial-Based Biopesticides: Methods and Protocols, vol 1477. Methods in Molecular Biology. Humana Press Inc, Totowa, pp 95–117.

Reil JB, Doorenweerd C, San Jose M, Sim SB, Geib SM, Rubinoff D (2018) Transpacific coalescent pathways of coconut rhinoceros beetle biotypes: Resistance to biological control catalyses resurgence of an old pest. Molecular Ecology 27:4459–4474.

Reil JB, San J, Rubinoff D (2016) Low variation in nuclear and mitochondrial DNA inhibits resolution of invasion pathways across the Pacific for the Coconut Rhinoceros Beetle (Scarabeidae: *Oryctes rhinoceros*). Proceedings of the Hawaiian Entomological Society 48: 57–69.

Richards NK, Glare TR, Aloali’i I, Jackson TA (1999) Primers for the detection of Oryctes virus from Scarabaeidae (Coleoptera). Molecular Ecology 8: 1552–1553.

Ridley AW, Hereward JP, Daglish GJ, Raghu S, McCulloch GA, Walter GH (2016) Flight of *Rhyzopertha dominica* (Coleoptera: Bostrichidae)-a spatio-temporal analysis with pheromone trapping and population genetics. Journal of Economic Entomology 109: 2561–2571.

Tsatsia F, Tsatsia H, Wratten H, Macfarlane B (2018) The status of Coconut Rhinoceros Beetle, *Oryctes rhinoceros* (L) Scarabaeidae: Dynastinae, in Solomon Islands. In: 51st Annual meeting of the society for invertebrate pathology, Gold Coast, Australia 2018. p 49

Unckless RL (2011) A DNA virus of *Drosophila.* PLoS One 6: e0026564

Young EC (1974) The epizootiology of two pathogens of the coconut palm rhinoceros beetle. Journal of Invertebrate Pathology 24: 82–92.

Zelazny B (1977a) Occurrence of the baculovirus disease of the coconut palm rhinoceros beetle in the Philippines and in Indonesia. FAO Plant Protection Bulletin 25: 73–77

Zelazny B (1977b) *Oryctes rhinoceros* populations and behavior influenced by a baculovirus. Journal of Invertebrate Pathology 29: 210–215.

Zelazny B, Alfiler AR (1991) Ecology of baculovirus-infected and healthy adults of Oryctes rhinoceros (Coleoptera: Scarabaeidae) on coconut palms in the Philippines. Ecological Entomology 16: 253–259.

Zelazny B, Alfiler AR, Lolong A (1989) Possibility of resistance to a baculovirus in populations of the coconut rhinoceros beetle (*Oryctes rhinoceros*) FAO Plant Protection Bulletin 37: 77–82.

Zelazny B, Lolong A, Pattang B (1992) *Oryctes rhinoceros* (Coleoptera: Scarabaeidae) populations suppressed by a baculovirus. Journal of Invertebrate Pathology 59: 61–68.

